# Primary Hyperparathyroidism Reveals Limited Adipose Remodeling in Human

**DOI:** 10.64898/2026.07.05.736588

**Authors:** Andrea Palermo, Fabio Zaccaria, Andrea Ninni, Anda Mihaela Naciu, Francesca Sciarretta, Luca Verteramo, Cecilia Gentile, Virginia Asia Barbetti, Lorenzo Nevi, Gaia Tabacco, Giorgia Conti, Federico Galli, Ciro Menale, Dario Tuccinardi, Filippo Longo, Pierfilippo Crucitti, Chiara Taffon, Anna Crescenzi, Katia Aquilano, Simone Carotti, Diego Sbardella, Daniele Lettieri-Barbato

## Abstract

**Background:** Preclinical models implicate the parathyroid hormone/parathyroid-hormone-related protein (PTH/PTHrP)-PTH1 receptor (PTH1R) axis in adipocyte lipolysis, adipose browning, and energy wasting. Whether this catabolic program is reproduced in vivo in humans remains unresolved. Primary hyperparathyroidism (PHPT), a condition of chronic endogenous PTH excess, provides a clinically relevant model to test the translational relevance of this pathway.

**Methods:** We combined population-scale analyses with a prospective human intervention study. PTH/PTH1R associations with body composition were evaluated in the UK Biobank and compared with PTH dynamics in cancer-associated cachexia using TRACERx proteomic data. In parallel, patients with PHPT were assessed before and after parathyroidectomy and compared with matched surgical controls. Biochemical parameters, circulating adipocytokines, DXA- and BIA-derived body composition, histology, UCP1 immunohistochemistry, and supraclavicular adipose tissue transcriptomic and proteomic profiles were integrated, with external validation in an independent supraclavicular adipose dataset.

**Results:** In the UK Biobank, apparent positive associations between circulating PTH/PTH1R signals and fat or lean mass were markedly attenuated after matching for age, sex, and BMI, arguing against a disease-specific adiposity effect of PHPT. In TRACERx, circulating PTH did not increase across BMI-adjusted weight-loss grades. In the prospective cohort, parathyroidectomy normalized PTH, calcium, and phosphate but did not induce coherent changes in glucose metabolism, lipid profile, inflammatory markers, body weight, fat mass, lean mass, or thermogenic adipose signatures. Supraclavicular adipose histology, UCP1 staining, RNA-seq, proteomics, pathway analysis, and external dataset reanalysis converged on the absence of browning or thermogenic activation. By contrast, PHPT was associated with a selective adipose-related secretory phenotype: adiponectin, adipsin, and retinol-binding protein 4 were reversible after surgery, whereas lipocalin- 2 and thrombospondin-1 remained elevated.

**Conclusions:** Chronic endogenous PTH excess is not sufficient to induce a detectable thermogenic or energy- dissipating adipose program in humans under basal clinical conditions. These findings challenge direct extrapolation from rodent PTH/PTHrP models and reposition the human PTH-adipose axis as a selective secretory and remodeling pathway rather than a dominant driver of adipose browning or wasting.

**Highlights:**

- PHPT provides an in vivo human model of chronic endogenous PTH excess.
- PTH/PTH1R associations with body composition are lost after stringent confounder control.
- Parathyroidectomy normalizes mineral metabolism without inducing adipose browning or wasting.
- Supraclavicular adipose histology, UCP1 staining, transcriptomics, and proteomics show no thermogenic activation.
- PHPT unmasks a selective adipose-related secretory signature with reversible and persistent components.

## INTRODUCTION

Parathyroid hormone (PTH) is classically recognized as a central regulator of calcium, phosphate, and skeletal homeostasis, but accumulating evidence indicates that the PTH/PTH-related protein (PTHrP)-PTH1 receptor (PTH1R) axis also intersects with adipose tissue biology and systemic energy metabolism (Izquierdo-Lahuerta, 2021; Tay Donovan and Bilezikian, 2024). Mechanistically, PTH can stimulate adipocyte lipolysis through cyclic adenosine monophosphate/protein kinase A (cAMP/PKA)-dependent phosphorylation of hormone- sensitive lipase and perilipin, supporting adipocytes as direct targets of PTH signaling (Larsson et al., 2016). Early ex vivo work showed that human PTH induces lipolysis in human adipose tissue (Taniguchi et al., 1987), and subsequent studies in primary human white adipocytes and human white/brown adipocyte models reported increased mitochondrial activity, oxidative respiratory capacity, thermogenic gene expression, hormone- sensitive lipase activity, and lipolysis following PTH or PTHrP receptor stimulation (Breining et al., 2021; Hedesan et al., 2019).

The biological relevance of this pathway has been most clearly demonstrated in experimental wasting states. Tumor-derived PTHrP promotes white adipose tissue browning, thermogenic gene expression, energy expenditure, and cachexia in mice, whereas PTHrP neutralization or adipocyte PTH1R disruption attenuates adipose browning and preserves tissue mass and function (Kir et al., 2016; Kir et al., 2014). Clinical observations have also linked circulating PTHrP to cancer-associated weight loss independently of hypercalcemia, inflammation, tumor burden, and performance status (Hong et al., 2016). Additional murine data suggest that PTHrP overexpression can remodel white and brown adipose depots and modulate diet- induced obesity, hepatic steatosis, and insulin resistance (Qin et al., 2022). Collectively, these studies define a biologically plausible PTH/PTHrP-PTH1R adipose program, but they also raise a critical translational question: does chronic endogenous PTH excess, in the absence of tumor-derived PTHrP, renal failure, or experimental sympathetic activation, induce comparable adipose remodeling in humans?

Primary hyperparathyroidism (PHPT), characterized by sustained endogenous PTH elevation, represents a stringent human model to address this question outside malignancy and chronic kidney disease (Walker and Bilezikian, 2000). A PHPT-focused translational study reported white adipose tissue browning, increased energy expenditure, reduced fat content, and lower body weight in PTH-overexpressing mice, together with an inverse association between PTH and body weight and greater brown/beige adipose activity in patients with PHPT (He et al., 2019). However, human clinical data remain heterogeneous. An earlier meta-analysis found that patients with PHPT were heavier than eucalcemic controls (Bolland et al., 2005), subsequent analyses suggested a nonlinear relationship between PTH and adiposity (Yuan et al., 2021), and matched clinical plus Mendelian-randomization analyses did not support a consistent association between PHPT or circulating PTH and anthropometric indices (Zhang et al., 2025). These discrepancies suggest that body weight and BMI alone may be insufficient to resolve whether chronic PTH exposure drives depot-specific lipolysis, thermogenic activation, or broader changes in body composition.

Here, we tested the hypothesis that chronic endogenous PTH excess remodels human adipose tissue by integrating population-level analyses with deep phenotyping of an interventional PHPT cohort. We first evaluated PTH/PTH1R relationships with body-composition traits in the UK Biobank and examined PTH behavior across weight-loss grades in the TRACERx cancer-cachexia cohort. We then profiled patients with PHPT before and after parathyroidectomy, combining biochemical assessment, adipocytokine measurements, DXA and BIA body composition, histology, UCP1 immunohistochemistry, and supraclavicular adipose tissue transcriptomics and proteomics. This design allowed us to distinguish a true thermogenic/wasting program from more restricted endocrine or remodeling responses and to define the human relevance of the PTH-adipose axis in vivo.

## RESULTS

### Population analyses do not support a robust PTH/PTH1R signal for human adiposity or wasting after confounder control

Chronic activation of the PTH/PTHrP-PTH1R axis has been linked to adipose lipolysis, browning, energy wasting, and body-weight regulation in preclinical models. To determine whether analogous signals are detectable at population scale in humans, we analyzed 24,861 UK Biobank participants, including 111 individuals with PHPT and 24,750 controls. As expected for PHPT, circulating PTH protein levels, reported as normalized protein expression (NPX), were markedly higher in affected participants than in controls (1.10 +/- 1.32 vs 0.03 +/- 0.84, p < 0.001). The PHPT group also differed in key demographic and anthropometric variables, being more often female (78.4% vs 51.7%), older (59.99 +/- 6.51 vs 56.01 +/- 8.04 years, p < 0.001), and having higher BMI (29.72 +/- 5.31 vs 27.32 +/- 4.67 kg/m2, p < 0.001) and total fat mass (30.81 +/- 11.33 vs 24.38 +/- 9.43 kg, p < 0.001). The circulating PTH1R-related proteomic signal was slightly higher in PHPT, but the effect size was negligible (0.07 +/- 0.31 vs 0.04 +/- 0.42, p = 0.023; Cohen’s d = 0.076), emphasizing the need for rigorous confounder control.

In multivariable linear models adjusted for age, sex, and disease status, higher z-scored PTH was associated with higher fat mass index (FMI; beta = 0.63, SE = 0.020, p < 0.001) and lean mass index (LMI; beta = 0.29, SE = 0.011, p < 0.001) (**Figure 1A, B**). PHPT status was also associated with higher FMI (beta = 0.74, SE = 0.294, p = 0.012) and LMI (beta = 0.40, SE = 0.165, p = 0.015), whereas PTH-by-group interactions were not significant, indicating no evidence that the PTH-body-composition relationship differed specifically in PHPT. PTH1R showed similarly modest positive associations with FMI (beta = 0.10, SE = 0.021, p < 0.001) and LMI (beta = 0.07, SE = 0.011, p < 0.001) without disease-specific interaction (**Figure 1C, D**). However, after propensity-score matching for age, sex, and BMI, these associations were substantially attenuated and no longer statistically significant. Thus, the apparent PTH/PTH1R relationships observed in the full cohort are best explained by baseline differences in demographic and body-composition structure rather than by a PHPT- specific adipose-remodeling effect.

**Figure 1.**
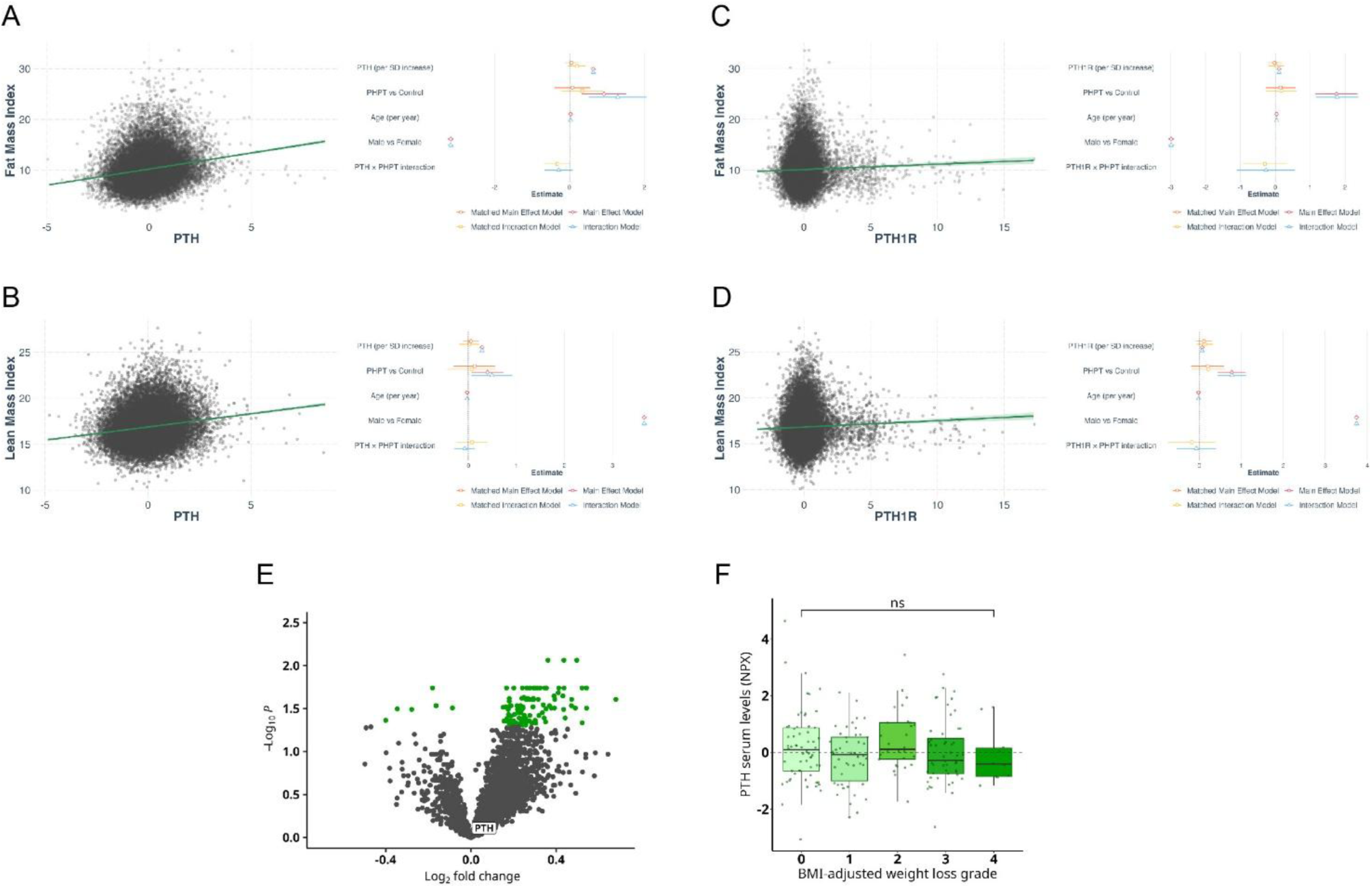
Population-scale analyses do not support a robust PTH/PTH1R signal for human adiposity or cachexia-associated wasting after confounder control. (A-B) Scatter plots showing the association between z-scored PTH and fat mass index (FMI; A) or lean mass index (LMI; B) in the UK Biobank analytic cohort, with accompanying regression summaries showing estimates from the main effect model and the interaction model. (C-D) Scatter plots showing the association between z-scored PTH1R and fat mass index (FMI; C) or lean mass index (LMI; D) in the UK Biobank analytic cohort, with accompanying regression summaries showing estimates from the main effect model and the interaction model. (E) Volcano plot representing the expression of PTH in patients with cancer-associated cachexia with significantly modulated proteins marked in green (padj < 0.05). (F) Circulating PTH levels across BMI-adjusted cancer-cachexia weight-loss grades (NS, not significant after multiple-testing correction).

We next asked whether circulating PTH tracks with human wasting in an independent cachexia setting. Using longitudinal plasma proteomics from the TRACERx non-small-cell lung cancer cohort, patients were stratified according to BMI-adjusted weight-loss grade. Circulating PTH did not increase across cachexia severity groups and did not correlate with the degree of BMI-adjusted weight loss (**Figure 1E, F**). Together with the UK Biobank analysis, these data argue against robust systemic activation of the PTH/PTH1R axis as a conserved driver of human adiposity loss or cancer-associated wasting.

### PHPT is not associated with altered body composition or basal metabolic phenotype in a matched clinical cohort

To directly test whether endogenous PTH excess alters human adipose metabolism in vivo, we recruited patients with PHPT undergoing parathyroidectomy and matched surgical controls undergoing surgery for benign euthyroid thyroid disease (**Figure 2A**). The design prioritized the clinical phenotype most represented in the UK Biobank PHPT group by focusing on older female participants and matching for major determinants of adipose metabolism, including body weight, BMI, thyroid function, 25-hydroxyvitamin D, and renal function (**Supplementary Figure 1A; Supplementary Table 2**). Supraclavicular adipose tissue was sampled because this human depot can harbor beige/brown adipocyte features and is therefore well suited to detect thermogenic remodeling (Cypess et al., 2009; Virtanen et al., 2009).

**Figure 2.**
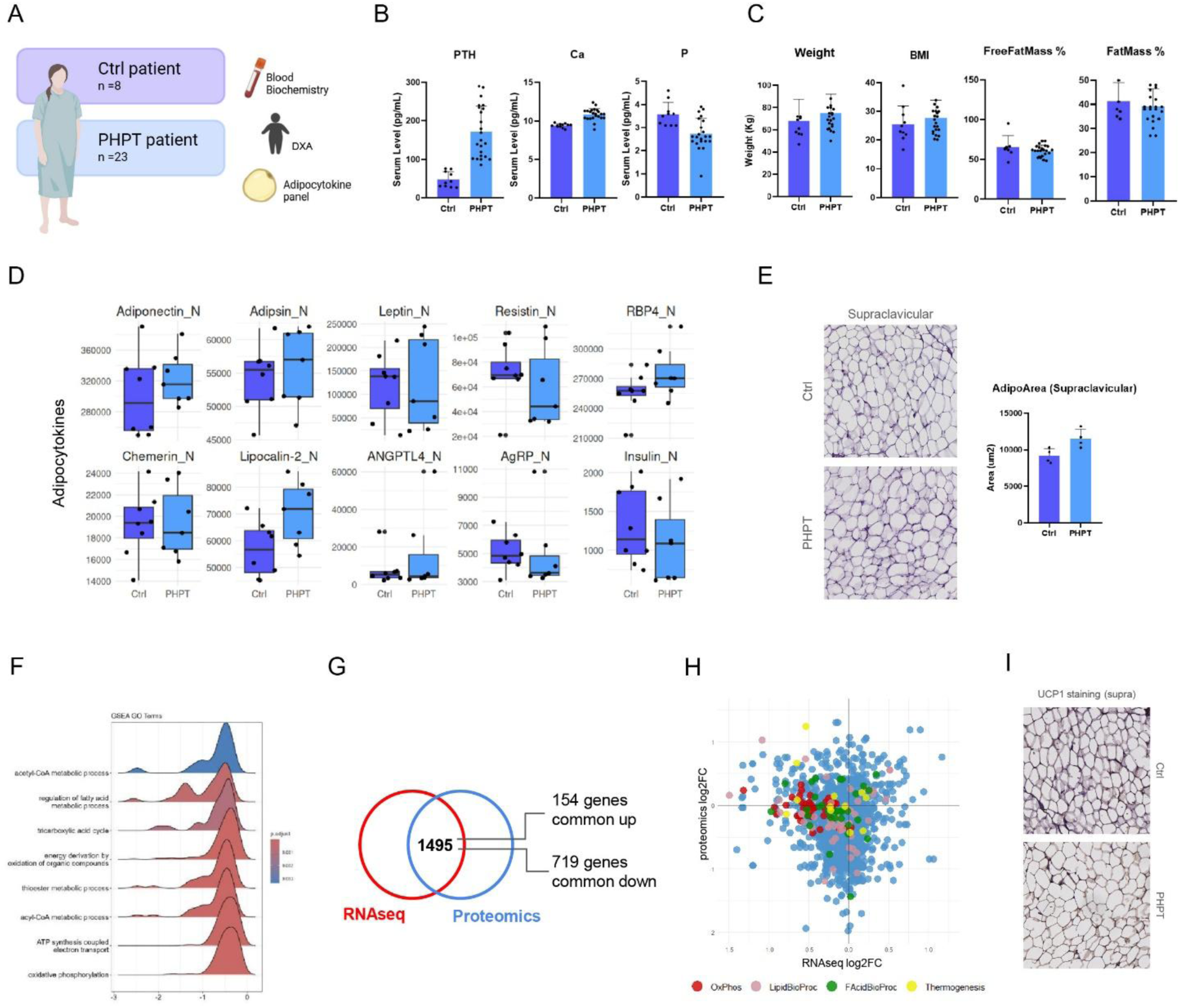
Molecular and phenotypic analyses reveal no thermogenic activation in adipose tissue from patients with PHPT. (A) Study design for the matched clinical cohort and adipose tissue sampling (cartoon created with BioRender). (B) Barplots plotting the mineral-metabolism profile of altered calcium-phosphate homeostasis in PHPT. (C) DXA/BIA-derived body-composition parameters showing no significant PHPT-associated adipose wasting. (D) Box-plots of circulating adipocytokine profile identifying a selective secretory signal rather than broad adipose dysfunction. (E) Hematoxylin and eosin staining and morphometric analysis of supraclavicular adipose tissue. (F) Transcriptomic Gene-Set Enrichment Analysis (GSEA) of statistically modulated pathways (padj < 0.05). (G) Transcriptome-proteome overlap analysis. (H) Two-dimensional transcriptome/proteome fold-change mapping of adipose metabolic and thermogenic pathways (oxidative phosphorylation, lipid biosynthetic process, fatty acid biosynthetic process and adaptive thermogenesis). (I) UCP1 immunohistochemistry showing absence of detectable UCP1-positive thermogenic adipocytes in supraclavicular adipose tissue.

Participants underwent fasting biochemical testing, DXA and BIA body-composition assessment, and circulating cytokine/adipocytokine profiling. As expected, PHPT was characterized by elevated PTH and calcium and reduced phosphate, whereas controls had normal mineral metabolism (**Figure 2B; Supplementary Figure 1A**). By contrast, glucose, triglycerides, HDL cholesterol, ALT, ESR, and CRP were comparable between groups. Total cholesterol and AST were modestly higher in PHPT, but the magnitude of these differences was small and did not define a coherent metabolic syndrome-like phenotype. Importantly, DXA and BIA showed no significant differences in body weight, BMI, fat mass, visceral adipose tissue, or appendicular lean mass between PHPT and controls (**Figure 2C**; **Table 1**). Exploratory cytokine profiling identified a selective endocrine signal rather than a broad inflammatory or adipose-mass phenotype: canonical adiposity markers such as leptin and adipsin were not consistently altered, whereas RBP4 and lipocalin-2 were increased and insulin was reduced in PHPT (**Figure 2D; Supplementary Figure 1B**).

**Table 1.**
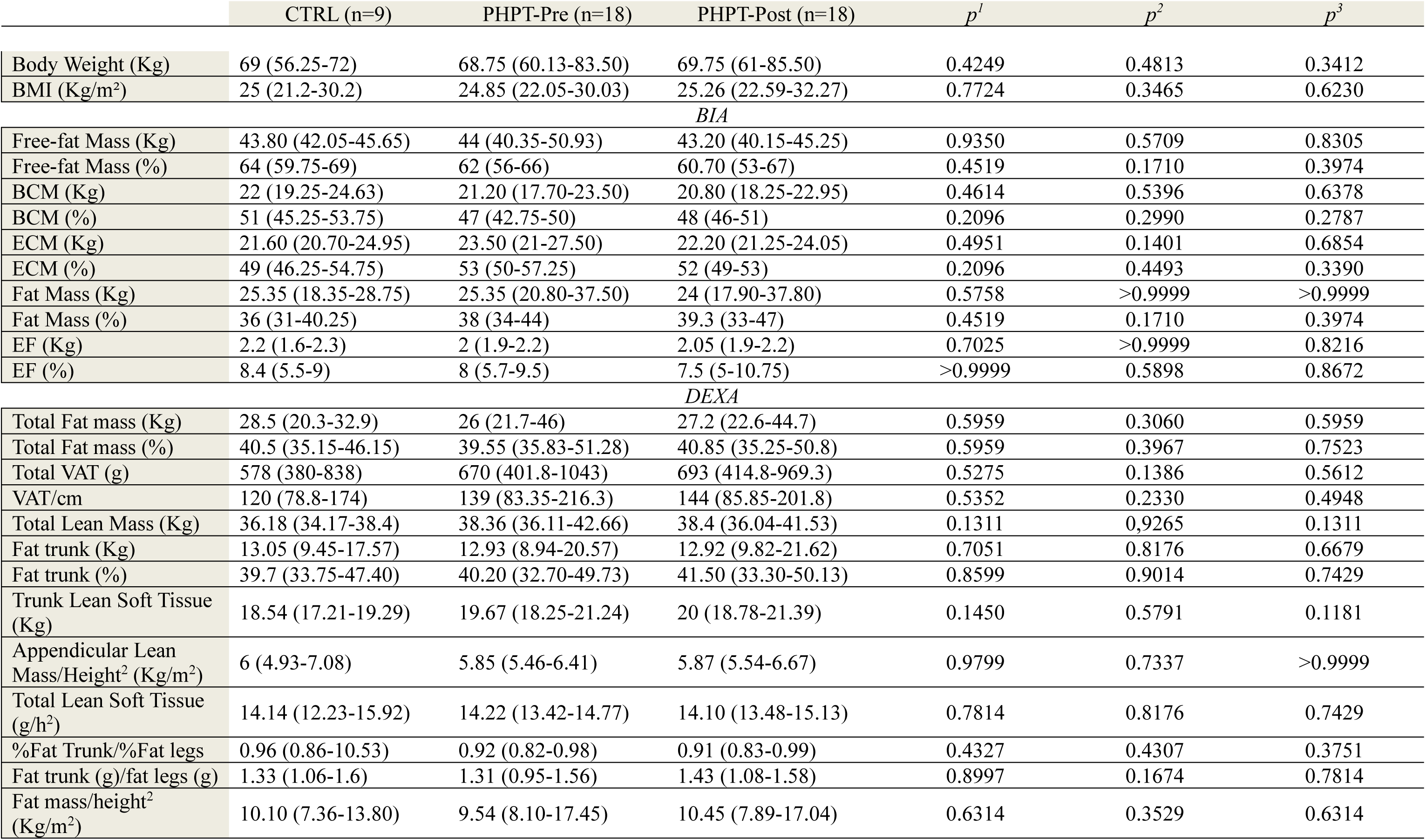
Body-composition parameters measured by bioelectrical impedance analysis (BIA) and DXA in control participants and PHPT patients before and after parathyroidectomy. Data are presented as median (25th-75th percentile). p^1^: CTRL vs PHPT-Pre (Mann-Whitney U test); p^2^: PHPT-Pre vs PHPT-Post (Wilcoxon signed-rank test); p^3^: CTRL vs PHPT-Post (Mann-Whitney U test).

### Supraclavicular adipose tissue from PHPT patients lacks histological and molecular evidence of thermogenic activation

Because whole-body body-composition measures may miss depot-specific adipose remodeling, we analyzed supraclavicular adipose tissue using histology, immunohistochemistry, transcriptomics, proteomics, pathway-level inference, and external dataset validation. Hematoxylin and eosin staining showed predominantly unilocular adipocytes in both PHPT and control samples, without the multilocular morphology expected for thermogenically activated brown/beige adipocytes (**Figure 2E**). Morphometric analysis did not support adipocyte shrinkage or lipid-droplet remodeling. Bulk RNA-seq and DIA proteomics were then performed in age- and BMI-matched PHPT and control samples from the same anatomical depot after standardized surgical sampling and overnight fasting. Principal-component analysis did not separate PHPT from controls in either omics layer (**Supplementary Figure 1C**). Differential analysis identified a restricted set of nominally modulated transcripts and proteins, but enrichment analyses did not reveal activation of thermogenesis, oxidative metabolism, or lipid-catabolic pathways (**Figure 2F**). Integrated transcriptome-proteome analysis further showed no coordinated upregulation of fatty-acid biosynthesis, oxidative phosphorylation, adaptive thermogenesis, or thermogenic effector genes (**Figure 2G, H; Supplementary Figure 1D**). Instead, the dominant directionality, when present, was neutral or reduced at the transcript level. Reanalysis of an independent supraclavicular adipose dataset from PHPT patients and controls (E-TABM-1119) confirmed the absence of a reproducible thermogenic gene-expression program (**Supplementary Figure 1E**). Finally, UCP1 immunoreactivity was undetectable in supraclavicular and jugular notch adipose tissue, and tissue architecture remained distinct from activated brown adipose tissue (**Figure 2I; Supplementary Figure 1F**).

Taken together, the histological, immunohistochemical, transcriptomic, proteomic, and external validation data converge on a negative result with mechanistic importance: despite sustained endogenous PTH excess, PHPT adipose tissue does not acquire an energy-dissipating, UCP1-positive, thermogenic phenotype under basal clinical conditions.

### Parathyroidectomy reveals a selective, partly reversible adipose-related secretory module

To determine whether correction of PTH excess modifies adipose-related endocrine signals, PHPT patients underwent longitudinal follow-up after parathyroidectomy (**Figure 3A**). Surgery effectively normalized PTH, calcium, and phosphate, confirming biochemical cure (**Figure 3B**). Renal and hepatic function, systemic inflammatory markers, glycemic variables, and lipid parameters remained broadly stable after surgery (**Supplementary Figure 2A**). Likewise, paired DXA and BIA analyses did not show significant postoperative changes in body weight, BMI, total fat mass, visceral adipose tissue, or lean mass (**Figure 3C**; **Table 1**), reinforcing the absence of macroscopic adipose wasting or rebound after PTH normalization. Circulating cytokine profiling nonetheless revealed a selective adipose-related secretory trajectory (**Figure 3D; Supplementary Figure 2B**). Adiponectin, adipsin, and RBP4 decreased after parathyroidectomy, identifying a reversible module linked to correction of the PHPT biochemical state. In contrast, lipocalin-2 and thrombospondin-1 remained elevated despite normalization of mineral metabolism, suggesting persistent inflammatory, matrix-related, or tissue-remodeling signals that are not rapidly reset by surgery (Sell and Eckel, 2007; Varma et al., 2008). This stratified behavior argues against a uniform PTH-driven adipose response and instead supports two components: a reversible secretory module and a more persistent remodeling/inflammatory module that appears at least partly PTH-independent.

**Figure 3.**
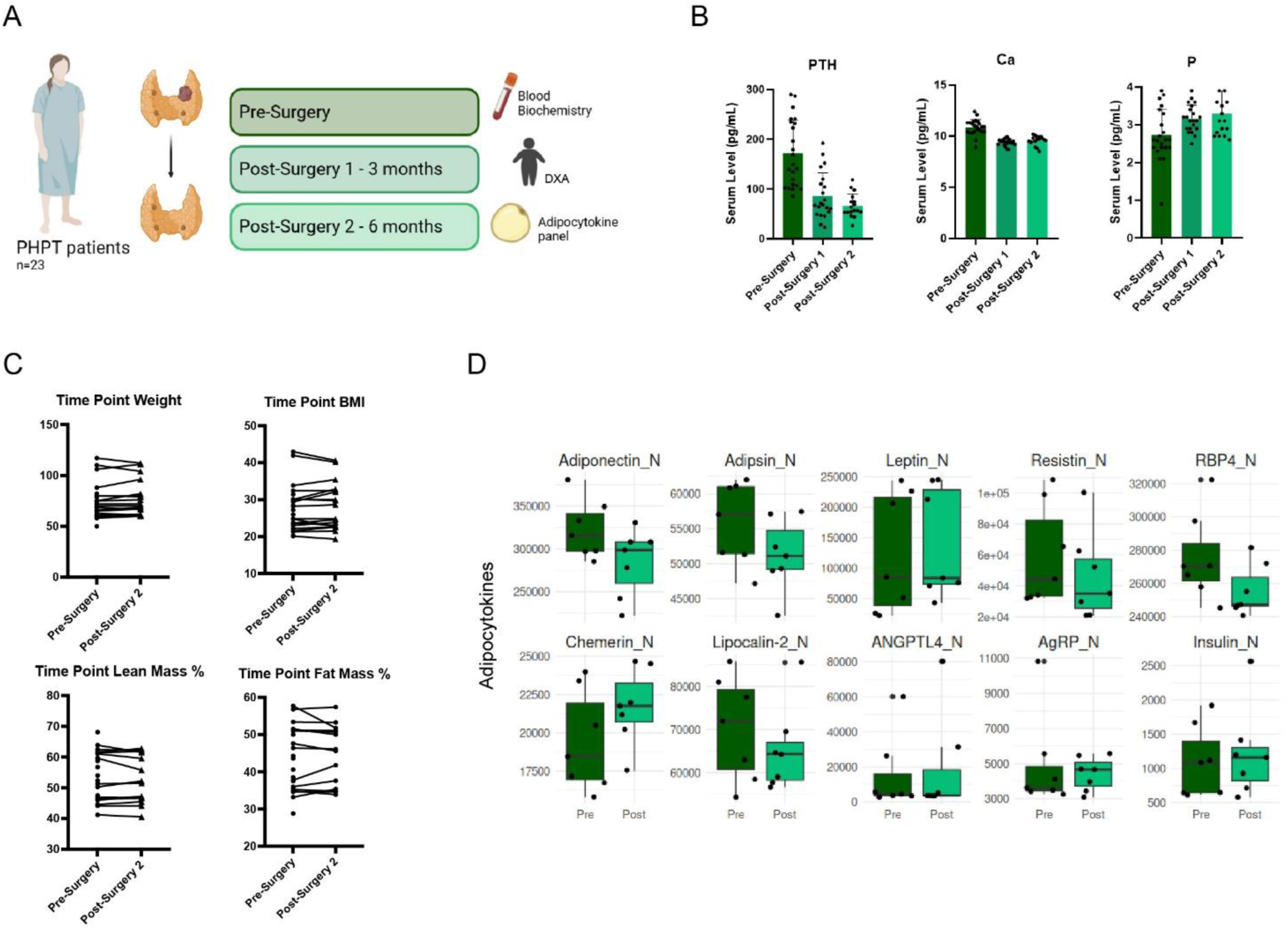
Parathyroidectomy reveals a selective, partly reversible adipose-related secretory module without body-composition remodeling. (A) Longitudinal study design before and after parathyroidectomy, cartoon created with BioRender. (B) Barplots showing the biochemical correction of PHPT after surgery in calcium-phosphate homeostasis. (C) Paired body weight, BMI, lean-mass, and fat-mass trajectories showing no coherent postoperative adipose wasting or rebound. (D) Box-plots of longitudinal adipocytokine profiling showing reversible changes in adiponectin, adipsin, and RBP4 and persistent elevation of lipocalin-2 and thrombospondin-1.

## DISCUSSION

Preclinical studies robustly support an adipose-catabolic program downstream of the PTH/PTHrP-PTH1R- cAMP/PKA axis. Tumor-derived PTHrP induces white adipose tissue browning, increases Ucp1/Pgc1a/Dio2 expression, elevates energy expenditure, and contributes to cachexia, while genetic or pharmacologic interruption of this pathway attenuates wasting (Kir *et al*., 2016; Kir *et al*., 2014). In adipocytes, PTH stimulates lipolysis through PKA-mediated phosphorylation of hormone-sensitive lipase (Larsson *et al*., 2016), and human ex vivo systems can respond to PTH/PTHrP with increased lipolysis and thermogenic gene expression (Breining *et al*., 2021; Hedesan *et al*., 2019). The central finding of the present study is that this biologically plausible program does not translate into a detectable thermogenic or wasting phenotype in vivo in patients with PHPT. Across population-level analyses, a prospective pre/post-parathyroidectomy cohort, matched surgical controls, supraclavicular adipose histology, UCP1 staining, transcriptomics, proteomics, and independent dataset validation, chronic endogenous PTH excess was not associated with adipose browning, UCP1-positive thermogenesis, fat-mass loss, or systemic wasting. These results help reconcile conflicting human and experimental observations. First, ligand and disease context are likely decisive. The strongest rodent phenotypes arise in PTHrP-secreting tumors or experimental systems with high local/paracrine signaling, additional inflammatory and catabolic mediators, and acute energy stress. PHPT instead represents chronic endocrine PTH excess without tumor-derived PTHrP and without the integrated cachexia program. Second, thermogenic competence is gated by sympathetic tone, ambient temperature, vascularization, innervation, and depot-specific cellular architecture. Mice are commonly housed below thermoneutrality, a condition that preactivates brown/beige adipose tissue and can amplify browning responses, whereas humans generally live closer to thermoneutral conditions (McKie and Wright, 2021). Third, adult human thermogenic depots are limited and heterogeneous, and recent single-cell and spatial atlases emphasize marked species, depot, progenitor, vascular, and immune-niche differences that can constrain endocrine responsiveness (Lazarescu et al., 2025; Massier et al., 2023). Within this framework, our supraclavicular samples did not show a coordinated thermogenic cell-state shift. The positive signal emerging from the study is not thermogenic but secretory. Parathyroidectomy selectively normalized adiponectin, adipsin, and RBP4, whereas lipocalin-2 and thrombospondin-1 remained elevated. This pattern is biologically informative because it separates a reversible adipose-related endocrine module from persistent inflammatory or extracellular-matrix-associated signals. RBP4 is linked to insulin resistance and adipose/liver metabolic dysfunction (Graham et al., 2006), adiponectin is a key anti-inflammatory and insulin-sensitizing adipokine (Tilg and Wolf, 2005), lipocalin-2 has been implicated in adipose inflammation and insulin resistance (Sell and Eckel, 2007), and thrombospondin-1 is associated with obesity, adipose inflammation, and extracellular-matrix remodeling (Varma *et al*., 2008). The absence of parallel changes in adipose mass, UCP1 expression, or thermogenic pathways indicates that these circulating changes should not be interpreted as evidence of browning; rather, they define a more restricted human PTH-adipose endocrine phenotype.

Several strengths increase the robustness of this interpretation. The study uses PHPT as a clinically grounded model of chronic endogenous PTH excess, incorporates a within-patient surgical intervention, includes matched controls, combines DXA and BIA body-composition assessment, interrogates a thermogenically competent human depot, and integrates histological, immunohistochemical, transcriptomic, proteomic, and independent public-dataset evidence. The conclusion is also strengthened by external human context: in TRACERx, PTH did not increase with BMI-adjusted weight-loss grade, arguing against a generalized PTH-driven cachexia signal in human cancer (Al-Sawaf et al., 2023). Importantly, the study does not rely on a single negative assay; multiple orthogonal readouts converge on the same conclusion.

Limitations should be acknowledged. The prospective cohort is modest in size and restricted to women undergoing surgery, which improves internal matching but limits generalizability to men, younger patients, and more severe or atypical PHPT phenotypes. Multi-omics was performed in a subset of participants, and bulk tissue profiling cannot resolve rare cell states or cell-type-specific receptor activation. The study assessed basal clinical conditions; cold exposure, beta-adrenergic stimulation, acute teriparatide challenge, PET-based thermogenic imaging, or receptor-centric functional assays may reveal conditional PTH responsiveness not captured here. Finally, longer postoperative follow-up may be required to determine whether persistent lipocalin-2 and thrombospondin-1 signals eventually normalize.

In conclusion, PHPT provides a rigorous human test of chronic endogenous PTH excess and shows that correction of mineral-hormone abnormalities is not accompanied by adipose browning, thermogenic activation, or systemic wasting. These findings challenge direct extrapolation from rodent PTH/PTHrP models to human adipose biology and define the human PTH-adipose axis as a selective secretory and remodeling pathway under basal clinical conditions rather than a dominant thermogenic effector.

## MATERIALS AND METHODS

### UK Biobank

UK Biobank is a prospective cohort study of more than 500,000 participants aged 38-73 years at recruitment (Sudlow et al., 2015). The initial sample included 503,386 individuals. Participants with cancer, chronic kidney disease, pregnancy, or vitamin D deficiency were excluded to reduce major confounding by conditions known to affect PTH, body composition, or systemic metabolism. The final analytic dataset included 24,861 participants, comprising 24,750 controls and 111 individuals with PHPT. Circulating PTH and the PTH1R- related proteomic signal were measured using Olink assays and reported as normalized protein expression (NPX) values on a log2 scale. For statistical analyses, biomarker values were standardized to z-scores to facilitate interpretation of regression coefficients. Body composition was assessed using fat mass index (FMI) and lean mass index (LMI), derived from total fat mass and lean mass measured by DXA. FMI and LMI were calculated as total fat mass or total lean mass divided by height squared (kg/m^2^).

Associations between PTH or PTH1R and body composition were assessed using multivariable linear regression models with FMI or LMI as continuous outcomes. Models were adjusted for age, sex, and disease group (PHPT vs controls). Interaction terms between each predictor and disease group were included in separate models to test whether associations differed between PHPT and control participants. Sensitivity analyses used propensity-score matching to generate balanced PHPT and control groups matched for age, sex, and BMI. The matching analysis was used to distinguish disease-specific effects from confounding by demographic and body-composition differences.

### Study population and design

This prospective observational study was conducted at the osteometabolic and thyroid disorders outpatient clinics of Fondazione Policlinico Universitario Campus Bio-Medico of Rome. Participants were consecutively recruited to minimize selection bias. The PHPT group included patients with a diagnosis of primary hyperparathyroidism who were candidates for parathyroidectomy according to current clinical criteria. Control participants had euthyroid benign nodular thyroid disease and were candidates for elective thyroid surgery; they were selected to match PHPT patients for age and BMI. General exclusion criteria for both groups were age less than 18 years, male sex, pregnancy or breastfeeding, severe hepatic disease, severe chronic kidney disease, medications known to affect calcium-phosphate or bone metabolism, drugs known to significantly influence body composition or fluid balance, and overt or subclinical hypo- or hyperthyroidism. Additional exclusion criteria for controls were any history or biochemical evidence of primary, secondary, or tertiary hyperparathyroidism and hypercalcemia or hypocalcemia. All participants provided written informed consent before inclusion. The study was conducted in accordance with the Declaration of Helsinki and was approved by the local Ethics Committee. Baseline evaluations included medical history, anthropometry, fasting blood sampling, BIA, and total-body DXA. PHPT patients were reassessed after surgery with biochemical testing, BIA, and DXA. Biochemical parameters, including calcium, phosphate, creatinine, 25-OH vitamin D, thyroid- stimulating hormone, total cholesterol, HDL cholesterol, triglycerides, glucose, AST/GOT, ALT/GPT, ESR, CRP, and PTH, were measured using automated routine laboratory methods.

### Bioelectrical Impedance Analysis and Dual-Energy X-ray Absorptiometry

Body composition was assessed using BIA. Measurements were performed with the participant in a fasting state and after voiding, following standard recommendations. Subjects were examined in the supine position with limbs slightly abducted to avoid skin contact. Fat mass, fat-free mass, and total body water were estimated using predictive equations provided by the manufacturer. The following variables were assessed: body weight (BW), BMI, fat-free mass (FFM), body cell mass (BCM), extracellular mass (ECM), fat mass (FM), and essential fat (EF). Fat-free mass, body cell mass, extracellular mass, and fat mass were estimated using predictive equations based on impedance measurements. Essential fat was derived from fat mass according to standard reference values. All measurements were conducted under standardized environmental conditions. DXA was used to assess total and regional body composition using a dual-energy X-ray absorptiometer (Hologic Discovery QDR Instrument, MA, USA). The analysis included appendicular lean mass adjusted for height (kg/m²), total lean mass adjusted for height (kg/m²), total fat mass adjusted for height (kg/m²), trunk fat mass, trunk fat percentage, trunk-to-leg fat ratio, and trunk fat-to-leg fat percentage ratio. Regional measurements were obtained for the trunk and appendicular compartments according to standard DXA segmentation. All scans were performed under standardized conditions following routine quality control procedures.

### Adipose tissue sampling

A subgroup of 7 PHPT subjects and 7 age- and BMI-matched controls underwent adipose tissue sampling based on availability and willingness to provide tissue. Participants completed an 8-hour fast before surgery. During the scheduled surgical procedure, and in accordance with standard clinical practice for parathyroidectomy or thyroidectomy, small adipose tissue samples were collected from the supraclavicular and jugular notch regions by the same trained physician. Samples were processed immediately after collection. One portion was flash frozen on dry ice for molecular analyses, and the remaining tissue was processed for histological or ancillary analyses.

### TRACERx

Plasma proteomics data were retrieved from the TRACERx study (Al-Sawaf *et al*., 2023). Data were analyzed using the OlinkAnalyze R package (v4.4.0). Proteomic quantification was represented as normalized protein expression (NPX) on a log2 scale. The analysis focused on PTH (OlinkID: OID30963). Patients were stratified into five groups according to BMI-adjusted weight-loss grade as defined by the original TRACERx authors: grade 0 (N = 64), grade 1 (N = 46), grade 2 (N = 29), grade 3 (N = 52), and grade 4 (N = 10). Differences in PTH abundance between weight-loss groups were assessed using two-sided Mann-Whitney U tests. Multiple comparisons were controlled using Benjamini-Hochberg correction.

### Bulk RNAseq

Total RNAs from cells and tissues were extracted using Direct-zolTM RNA MiniPrep (ZYMO RESEARCH) according to the manufacturer’s instructions. Total RNA was quantified using the Qubit 4.0 fluorimetric Assay (Thermo Fisher Scientific). Libraries were prepared from 50 ng of total RNA using the NEGEDIA Digital mRNA-seq research grade sequencing service (Next Generation Diagnostic srl), which included library preparation, quality assessment and sequencing on a NextSeq 500 sequencing system using a paired-end, 75 cycle strategy (Illumina Inc.).

The bulk RNAseq data were quality-checked using FastQC v0.11.9 and subsequently aligned to the GENCODE Human Release 47 (GRCh38p14) reference genome using HISAT2 v2.2.1 with default parameters. The number of reads for all RefSeq genes was counted using featureCounts v2.0.6 enforcing the multi-mapping option. The resulting count matrix was analyzed in R v4.6.0 using DESeq2 v1.48.2. Differential expression was used as an exploratory discovery layer because of the limited number of human adipose samples; nominal p < 0.05 was used to define candidate transcripts, whereas pathway-level interpretation emphasized directionality, concordance with proteomic data, and external validation rather than isolated gene-level significance. Enrichment analyses were performed with clusterProfiler v4.16.0. Gene-set enrichment analysis (GSEA) used the ranked expression matrix, and over-representation analysis (ORA) was performed on the most modulated genes. Most of the results were plotted using ggplot2 v4.0.3, the heatmaps were created with pheatmap v1.0.13.

### Microscopy analysis

Supraclavicular and jugular notch adipose specimens were stained with hematoxylin and eosin (H&E) according to standard protocols. Sections of 3 µm thickness were prepared for immunohistochemistry (IHC). Endogenous peroxidase activity was blocked by incubating sections in 3% hydrogen peroxide for 30 minutes. Antigen retrieval was performed at 95 °C for 40 minutes using Citrate buffer 10X pH 6.0 (Sigma-Aldrich, Cat. #C9999-100ML) diluted at 1X with distilled water. The sections were then incubated overnight at 4 °C in a humidified chamber with a primary antibody against UCP1 (ABCAM, Cat. #Ab10983), diluted 1:1000 in 5% BSA/PBS. Detection of the primary antibody was performed using the MACH 4 Universal HRP Polymer (BioCare Medical, Cat. #M4U534L) according to the manufacturer’s instructions. Visualization was achieved using 3,3′-diaminobenzidine as the chromogen (Dako, Cat. #K3468), and nuclei were counterstained with hematoxylin. Sections were scanned using Hamamatsu NanoZoomer 2.0 RS and five fields were randomly chosen for the statistical analysis (magnification x100) for each sample.

### Proteomic Analysis

#### Sample Preparation

LC/MS-grade water was used throughout the procedures for protein extraction, buffer preparation, and mass spectrometry analysis. Protein concentration in lysed samples was determined by a bicinchoninic acid assay (BCA) following the manufacturer’s protocol. For each experimental condition, a total of 30 µg of protein was precipitated from lysed samples by diluting the lysates with a 4-fold volume of pure acetone kept overnight at -20 °C. The following day, proteins were pelleted by centrifugation (13,000 × g, 10 min, 4 °C). The protein pellet was then washed once with 80% cold acetone and pelleted again by centrifugation as indicated above. Proteins were resuspended in 9 M urea buffer (50 mM Tris-HCl, pH 8) and sonicated in a water bath for complete solubilization and denaturation. Thereafter, reduction (5 mM dithiothreitol, 45 min, room temperature) and alkylation (10 mM iodoacetamide, 30 min in the dark) were performed. Protein digestion was carried out using mass spectrometry-grade trypsin (Merck Millipore, Darmstadt, Germany) at a 1:50 trypsin-to-protein ratio, after diluting the urea concentration to 1 M using 50 mM Tris-HCl (pH 8). Digestion was allowed to proceed overnight at 28 °C. The reaction was quenched with 0.4% trifluoroacetic acid, and peptides were desalted and cleaned using C18 stage-tips (Fisher Scientific, Waltham, MA) according to the manufacturer’s instructions. Eluted peptides were dried in a SpeedVac concentrator (Labconco, Kansas City, MO) and resuspended in 5% acetonitrile (ACN), 0.1% formic acid (hereafter referred to as Solvent A). Pooled GPF fraction was prepared by mixing 1 µL from each tryptic sample included in the study. Six consecutive injections (GPF fractions 1–6) were performed at the midpoint of the sequence. Instrument performance was routinely verified by injecting a standard HeLa digest (Thermo Fisher Scientific, Waltham, MA, USA) every 36 hours of continuous analysis.

#### LC/MS-MS Settings

Chromatographic separations were performed on an Ultimate 3000 system (Thermo Fisher Scientific, Waltham, MA, USA). The column oven temperature was maintained at 45.0 °C, while the autosampler was set to 4.0 °C. Samples (1 µg of peptides each) dissolved in 100% H_2_0 with 0.1% formic acid (FA) were loaded using the loading pump at a flow rate of 20.0 µL/min, and analytical separation was carried out by the nano- capillary (NC) pump at a constant flow rate of 0.300 µL/min. A 60-minute gradient was employed as follows: an initial 2-minute isocratic step at 6% solvent B (80% acetonitrile, 0.1% FA), a linear increase from 6% to 31% B over 40 minutes, a ramp to 50% B by minute 46, a column wash at 99% B until minute 54.5, and a final re-equilibration at 6% B until minute 60. Mass spectrometry data were acquired in DIA mode on an Orbitrap Exploris 240 system (Thermo Fisher Scientific, Waltham, MA, USA). The NSI source was operated with a transfer tube temperature of 280 °C, with the spray voltage set to 1500 V. For MS1 scans, the sample method utilized an Orbitrap resolution of 120,000, an AGC target of 300%, and “Auto” maximum injection time, covering a mass range of m/z 395–1205. In contrast, GPF methods were acquired at a resolution of 60,000 with a 40% AGC target and a 60 ms maximum injection time; the total mass range was divided into six staggered 110 m/z fractions (GPF1: 395–505; GPF2: 495–605; GPF3: 595–705; GPF4: 695–805; GPF5: 795–905; GPF6: 895–1005). For MS1 and MS2 DIA scans, both strategies utilized an Orbitrap resolution of 30,000. The MS2 sample method featured a 21 m/z isolation window, 28% HCD collision energy, a 3000% AGC target, and “Auto” injection time, with data collected in Profile mode. The GPF methods utilized 4 m/z isolation windows with a 1 m/z overlap, 33% HCD energy, a 400% AGC target, and a 60 ms maximum injection time, with data acquired in Centroid mode

#### LC-MS/MS Data Processing and Database Searching

Raw DIA data were processed using DIA-NN software (v2.2.0). Initially, a pilot search in library-free mode was performed against a human FASTA database (containing canonical and reviewed sequences), including a list of common contaminants. Default parameters were utilized, except that the maximum charge state was increased to 5 and one missed cleavage was allowed, in order to determine the optimal MS1 accuracy and window settings.

All raw files, along with the gas-phase fractionation (GPF) files, were then processed using these optimized parameters (MS1 accuracy: 5 ppm; MS2 accuracy: 10 ppm; window radius: 7) to generate a GPF-refined library, with match-between-runs set to “Off”. For database searching, carbamidomethylation of cysteine was set as a static modification, while oxidation of methionine and N-terminal acetylation were set as variable modifications. N-terminal methionine excision was enabled. A maximum of two variable modifications per peptide and one missed cleavage were allowed, using Trypsin/P enzymatic specificity.

Finally, all experimental raw DIA files (excluding GPF files) were searched against the GPF-refined library using the same parameters. False discovery rate (FDR) assessment for both precursors and proteins was controlled at a *q*-value less than 0.01 using the target-decoy validation strategy implemented in DIA-NN.

#### Downstream Data Analysis and Statistics

All downstream proteomic and statistical analyses were performed in RStudio (v4.5). Data manipulation, cleaning, and preprocessing were carried out using the tidyverse suite (including dplyr, tidyr, stringr, tibble, and stringi), along with gtools, arrow, readxl, and writexl.

Contaminants were removed, and protein identifications were filtered to retain only those with at least one proteotypic peptide. Protein intensities were subsequently normalized via median centering. Proteins were further filtered to retain only those identified in at least 5 out of 7 samples per experimental group. Missing values were imputed using a Gaussian shift of 1.8 and a standard deviation (SD) of 0.3 (Perseus-like imputation); however, imputed values were strictly utilized for Principal Component Analysis (PCA), which was performed using factoextra.

Differential protein abundance analysis was conducted on non-imputed data using a moderated Bayesian t-test implemented in the limma package. Primary significance thresholds were defined as absolute log2 fold-change >= 1 and Benjamini-Hochberg-adjusted p value <= 0.05. For exploratory clustering and biological rationalization, proteins with absolute log2 fold-change >= 1 and unadjusted p value <= 0.05 were also examined. Proteins identified as significantly upregulated or downregulated were submitted to Gene Ontology enrichment analysis across Biological Process, Molecular Function, and Cellular Component categories. Functional enrichment and pathway analysis were performed with clusterProfiler using org.Hs.eg.db and msigdbr databases. Enriched terms were filtered using a Benjamini-Hochberg false discovery rate <= 0.05.

Proteins identified as significantly upregulated or downregulated between the experimental groups were submitted to Gene Ontology (GO) enrichment analysis across Biological Processes (BP), Molecular Function (MF), and Cellular Component (CC) categories. Functional enrichment and pathway analysis were performed via clusterProfiler, utilizing the org.Hs.eg.db and msigdbr databases. Protein annotations were retrieved using UniprotR. Data visualizations were generated using ggplot2, ggrepel, pheatmap, plotly, tidytext, and enrichplot, while overlap representations were constructed using VennDiagram, eulerr, and ggVennDiagram. Enriched terms and pathways were filtered using a BH-corrected False Discovery Rate (FDR) ≤0.05

## DATA AVAILABILITY

The mass spectrometry proteomics data have been deposited in the ProteomeXchange Consortium via the PRIDE partner repository (Perez-Riverol et al., 2025).

Processed transcriptomic matrices and analysis scripts are available from the corresponding author upon reasonable request; deposition of raw sequencing files should be completed before final acceptance if required by the journal.

## ACKNOWLEDGEMENTS

This work was supported by the Associazione Italiana per la Ricerca sul Cancro (AIRC) under MFAG 2023 – ID. 28842 project to D.L.-B.

The authors are grateful to the Italian Ministry of Health and Fondazione Roma for the support.

This research has been conducted using the UK Biobank Resource under Application Number 1090583.

## AUTHOR CONTRIBUTIONS STATEMENT

A.P. conceived the study, designed the analyses, acquired the data, interpreted the results and wrote the manuscript. A.N.M. and G.T. conceived the study, designed the analysis, interpreted the results and contributed to the text. F.Z., A.N., L.V., C.G. and G.C. performed bioinformatic analyses and interpreted the results. F.S. processed samples for the experimental part and contributed to formal analysis. D.T., P.C., F.L., A.C., and C.T. acquired the data, interpreted the results and contributed to the text. V.A.B. and F.G. processed samples for the experimental part. C.M. designed the analysis, interpreted the results and contributed to text. L.N. and S.C. performed hematoxylin-eosin and immunohistochemistry analyses. D.S. performed proteomics analysis and contributed to the text. F.Z., A.N., C.G., K.A. and D.L.-B. wrote the manuscript and edited the figures.

## COMPETING INTERESTS STATEMENT

Authors declare no competing interest.

---

**Supplementary Figure 1.**
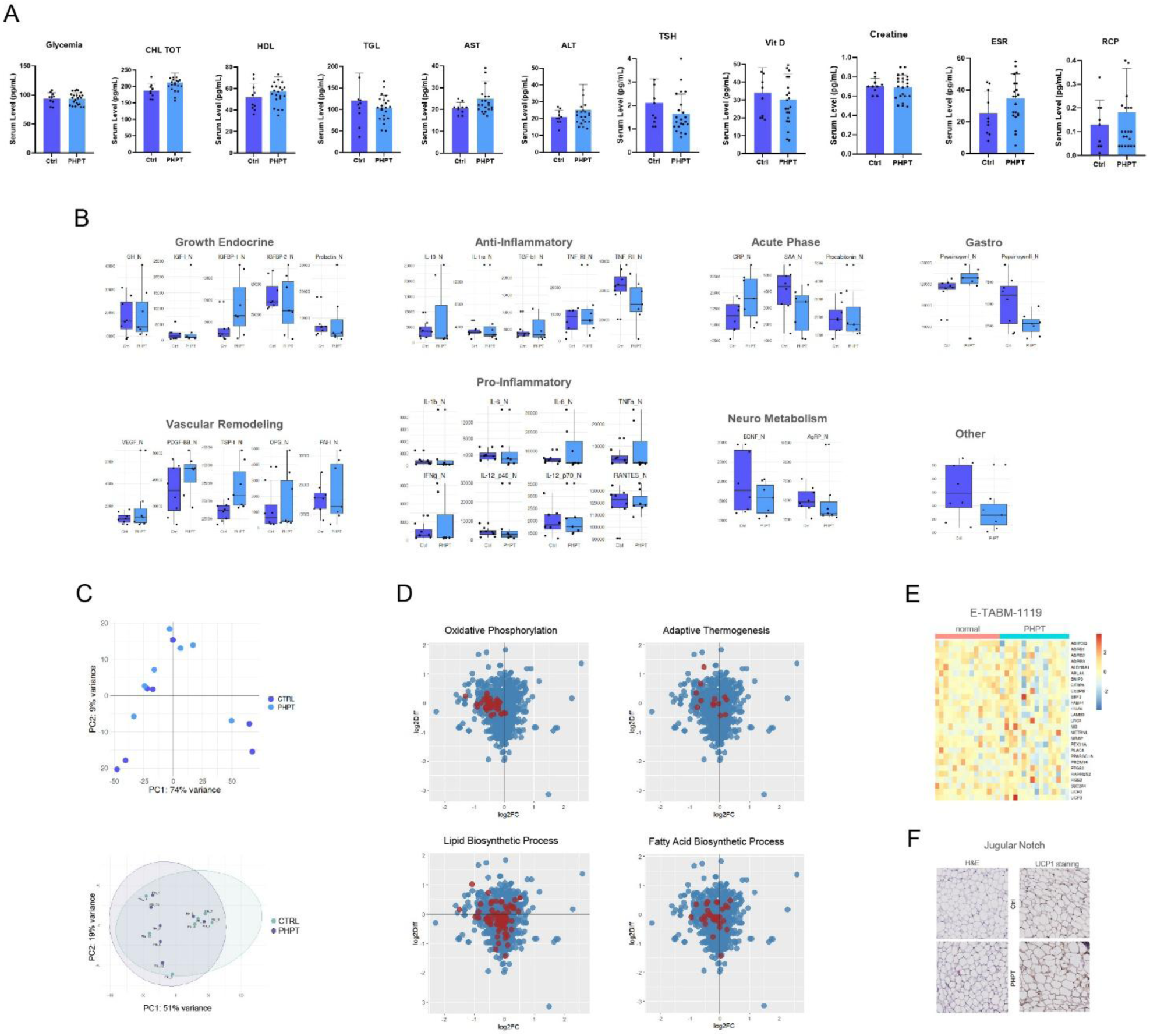
Baseline biochemical, cytokine, histological, and multi-omics quality-control analyses supporting the absence of PHPT-associated adipose thermogenic activation.

**Supplementary Figure 2.**
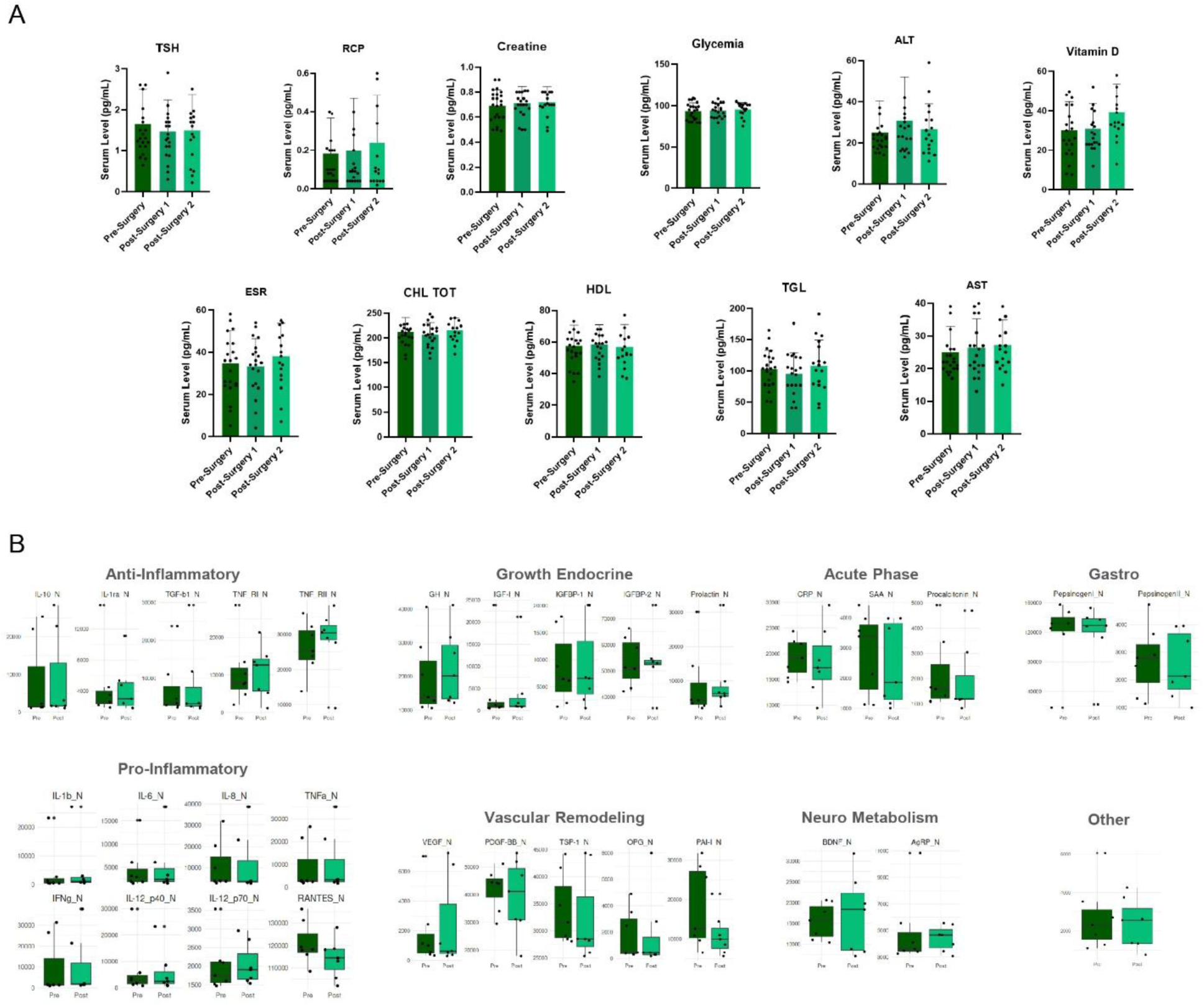
Longitudinal biochemical and cytokine analyses after parathyroidectomy.

**Supplementary Table 2.**
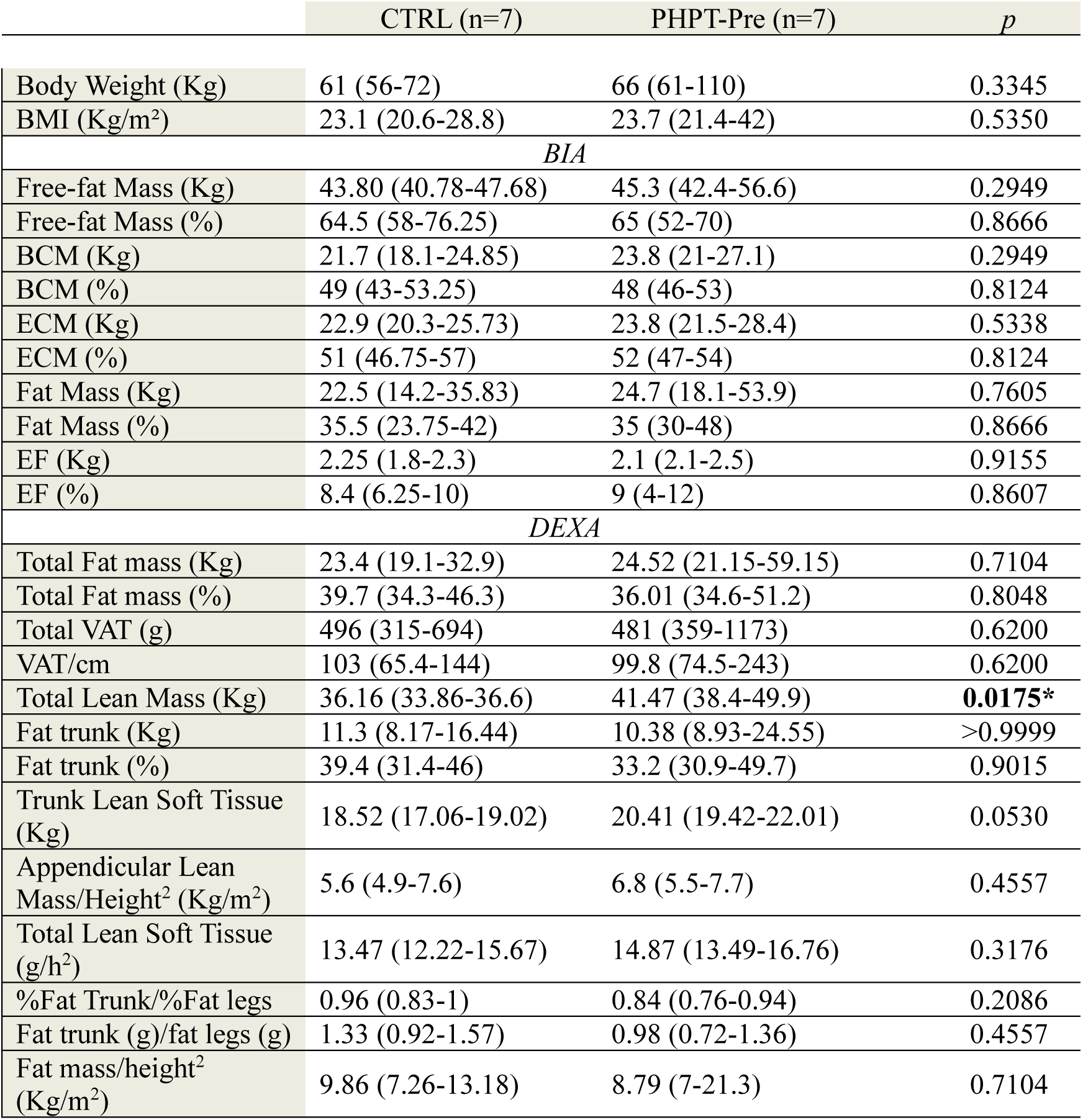
Body-composition parameters measured by bioelectrical impedance analysis (BIA) and DXA in control participants and PHPT patients before and after parathyroidectomy. Data are presented as median (25th-75th percentile). p^1^: CTRL vs PHPT-Pre (Mann-Whitney U test); p^2^: PHPT-Pre vs PHPT-Post (Wilcoxon signed-rank test); p^3^: CTRL vs PHPT-Post (Mann-Whitney U test).

## Notes

### Competing Interest Statement

The authors have declared no competing interest.

